# Seston fatty acids to quantify phytoplankton functional groups and their spatiotemporal dynamics in a highly turbid estuary

**DOI:** 10.1101/607614

**Authors:** José-Pedro Cañavate, Stefanie van Bergeijk, Enrique González-Ortegón, César Vílas

## Abstract

Phytoplankton community composition expresses estuarine functionality and its assessment can be improved by implementing novel quantitative fatty acid based procedures. Fatty acids have similar potential to pigments for quantifying phytoplankton functional groups but have been far less applied. A recently created dataset containing vast information on fatty acids of phytoplankton taxonomic groups was used as reference to quantify phytoplankton functional groups in the yet undescribed Guadalquivir River Estuary. Twelve phytoplankton groups were quantitatively distinguished by iterative matrix factor analysis of seston fatty acid signatures in this turbid estuary. Those phytoplankton groups including species unfeasible for microscopy identification (coccoid or microflagellated cells) could be quantified when using fatty acids. Conducting monthly matrix factor analyses over a period of two years and the full salinity range of the estuary indicated that diatoms dominated about half of the phytoplankton community spatiotemporally. The abundance of Cyanobacteria and Chlorophytes was inversely related to salinity and little affected by seasonality. Euglenophytes were also more abundant at lower salinity, increasing their presence in autumn-winter. Coccolithophores and Dinophytes contributed more to phytoplankton community at higher salinity and remained little affected by seasonality. Multivariate canonical analysis indicated that the structure of the estuarine phytoplankton community was most influenced by salinity, secondly influenced by water temperature, irradiance and river flow, and unaffected by nutrients. Fatty acids are especially suited for phytoplankton community research in high turbid estuaries and generate outcomes in synergy with those derived from classical pigment assessments.

## Introduction

Estuaries are complex and variable ecosystems in which internal processes are strongly dependent on highly fluctuating environmental factors expressed in spatial and temporal scales [1]. Understanding mechanisms underlying the functioning of estuaries is not easy [2] and it can be even more complicated when anthropogenic impacts are included. Within the complex estuarine structure, phytoplankton community is principally modelled by factors such as temperature, salinity, irradiance, nutrients, turbidity, turbulence and water residence time [3,4]. Phytoplankton community composition is a main indicator of the status and functionality of aquatic ecosystems. Hence, advancing methods to more precisely determine phytoplankton community structure will help to a better understanding on what phytoplankton can actually explain in estuaries. It is also evident that consequences of anthropogenic impacts (eutrophication, dredging, damming) would be better understood if more detailed information on phytoplankton community structure is available.

Phytoplankton community structure in most aquatic ecosystems has been traditionally studied from microscopic cell counting. While light microscopy allows high specific taxonomic resolution for the skilled analyst, it is a time-consuming task and has some limitations due to precluding identification of many picoplankton and nanoplankton species. It also fails in identifying those cells that are fragile and can be either distorted or disrupted by the fixatives used in microscopy [5]. To compensate for these limitations, chemotaxonomy based on the characteristic pigment composition of phytoplankton groups appeared as an alternative and complementary method in phytoplankton community research [6]. Inferring the relative abundance of phytoplankton taxonomic groups from their pigment signatures is a common practise that has led to important advances in phytoplankton ecology. However, this method still shows some inaccuracies [7] and it is partially restricted by the absence of specific signature pigments in some important taxa, as well as the occurrence of common pigments being shared by different taxonomic classes [8]. The high pigment content variability induced by the effects of light and nutrients [9] represents an additional limitation of the method. Fatty acids (FAs) are also selective biomarkers of specific phytoplankton groups [10,11] and can be used in a similar fashion as pigments to quantitatively infer the structure of microbial assemblages. Light-induced variability on phytoplankton FAs is lower than that of pigments [12] and other external factors also exert a noticeably lower influence on FAs than taxonomy [10], representing an interesting strength for the use of FAs as chemo markers. On the other hand, phytoplankton group-specific FAs can also be found in other non-phototrophic particulate material and that makes FAs to be less selective than pigments to specifically identify primary producers. While bacteria can be clearly differentiated from phytoplankton on the basis of their FAs [11], some overlapping might occur between terrestrial detrital plant material (depending on their degradation stage) and some phytoplankton groups belonging to the chlorophyte. Using FAs to assess aquatic microbial community structure implicitly computes some heterotrophic protist groups that are not detected by pigments and for which there are no current reference FAs signatures. FAs have been far less used than pigments to infer structure of phytoplankton assemblages and from the few studies available [12,13] FAs were found to generate results in synergy with those obtained from pigment analysis. There is thus scope to advance applying FAs use in phytoplankton ecology focusing in their contribution to improve quantitative taxonomic definition of phytoplankton assemblages.

The scarce use of FAs as quantitative phytoplankton biomarkers is related with the insufficient availability of those adequate reference libraries that are needed to carry out the CHEMTAX iterative methodology normally followed in this kind of studies [14]. Among the paucity of works using of FAs to infer phytoplankton community structure in aquatic environments, only a few of them [12, 15–17] can be highlighted for estuarine waters. In general, most FAs-based studies in aquatic ecosystems have been restricted to the qualitative estimation of a handful of phytoplankton groups abundance due to the already commented lack of FAs reference libraries. In pioneering works [12, 18, 19], site-specific reference FAs libraries had to be produced from locally isolated and cultured microorganisms as a previous step to produce the first quantitative results on phytoplankton community structure. Performing isolation, culturing and fatty acid analysis of those microalgae representatives of the studied phytoplankton community is a complex task that may demand even longer time than that required by the specific study itself. It has also the uncertainty as to whether the complete set of taxa present in the studied place is covered. To avoid such a tedious task, an updated reference fatty acid library compiling updated information for 19 phytoplankton classes has been recently generated [11]. The aim of this library is providing a new resource applicable anywhere to quantify phytoplankton community structure, backed by the finding that taxonomy accounts for four times more variation than the most influencing external factors in phytoplankton FAs [10].

The fatty acid approach can be of special interest to infer phytoplankton community structure in aquatic environments exhibiting particularly difficult conditions for microscopy cell identification, as can be those found in turbid estuarine waters. In this regard, the Guadalquivir River Estuary (GRE), located in SW Spain, has been characterized as a highly turbid estuary due to excessive inert sediment concentration [20]. It is interesting to note that no specific study on the phytoplankton community in the GRE has been published to date, whereas in the nearby less turbid Guadiana Estuary studies on phytoplankton are frequent [20, 21]. Being aware of the technical difficulty for phytoplankton research in the GRE, present study had the main goal of using FAs as biomarkers for the quantitative determination of the phytoplankton community structure in the GRE, as well as its dynamic under environmental forcing. The work had the following specific objectives: i) Demonstrate the suitability of iterative analytical methodology using the largest known to date FAs reference library to quantify contribution of phytoplankton taxonomic groups to community in estuarine highly turbid waters; ii) Use seston FAs signatures to describe the seasonal dynamics of the main phytoplankton groups along the salinity gradient in the GRE; iii) Identify the influence of environmental factors on the structure of phytoplankton community in the GRE.

## Materials and methods

### Studied area and sample collection

The GRE comprises the last 110 km of the 680 km length Guadalquivir River, draining a basin of 63,822 km^2^ [20]. It is a strongly hydrologically regulated river, with 57 dams along its course, and the estuary section between the river mouth and the port of Seville has been converted into an 84 km navigating channel (50 km shorter than the original course). The GRE keeps a minimum depth of 6.5 m by means of periodic dredging and it is completely isolated from the fringing marshes. As a result of this strong regulation, freshwater inputs to the estuary have decreased an average 60%, with extreme reduction in dry years [20]. The Alcala dam is the last hydrological regulation point and sets the upper limit of the estuary, 110 km upstream the river mouth. The estuary has a cross-section of 580 m^2^ at the Alcala dam and 4525 m^2^ near the mouth [23]. Freshwater discharge from the Alcala dam represents around 80% of the river water input to the estuary and the relatively low water volume dammed minimises its regulation capacity during sporadic stormy events of high rainfall. Low river flow regime (below 40 m^3^ s^−1^) occurs over 75% of the year and during this period the estuary is well mixed and tidally dominated [23]. Occasional dam discharges above 400 m^3^ s^−1^ turns the estuary into a completely fluvial-dominated hydrological regime. Tidal amplitude is around 3 m, situating the GRE in the meso-tidal category.

Water samples from the GRE were taken between November 2013 and October 2015 every new moon, accounting for a total of 25 consecutive sampling times. In each sampling, near-surface (20 cm depth) water was collected from an anchored boat during both flood and ebb tide at two different points that were 32 km (Tarfía) and 8 km (Bonanza) distant from the river mouth, respectively. This sampling covered the full salinity range of the estuary. On board samples were sieved through a 100 µm mesh, stored in 25 l containers and sent within 24 h to the IFAPA Centro El Toruño laboratory for phytoplankton identification and particulate matter fatty acid analysis. Samples were arranged considering season (winter, spring, summer, autumn) and estuary salinity range (levels 0-5, 5-10, 10-20 and 20-35 ppt) as crossed fixed factors.

### Analytical procedure

Water aliquots (25 ml) were taken for taxonomic identification of microalgae species under the microscope. Samples were directly observed at 200× and 400× magnification without the use of any fixative. Species were enumerated without any sample settlement since, in such case, high inorganic particle content prevented microscopic cell identification. Observation fields were extended until no new taxon could be detected in the sample and results were then expressed as percent cell composition. Microscopically detected microalgae were identified at the genus level and their relative cell abundance was seasonally compiled for each of the four salinity ranges established in the estuary. Identifying microalgae at the genus rank allowed creating phytoplankton functional groups at the class level needed for comparison with results from the inferring models.

Due to excess of suspended inert material, particle retention in glass fiber filters was inefficient to ascertain adequate sample size for lipid extraction. To overcome such inconvenience, a total of 25 l water sample was centrifuged using a continuous centrifuge RINA SRP 2C (Riera Nadeu SA, Barcelona, Spain) equipped with a 0.2 l clarifying capacity rotor. The centrifuge was operated at 7000 g with an estuary water sample constant inflow of 90 l h^−1^. After centrifugation of every 25 l water sample, the centrifuge was stopped and the rotor detached to enable collecting the centrifuged particles from a Teflon sheet covering the inner wall of the rotor. The centrifuged material was easily scraped with a spatula, lyophilized and transferred to storage bottles that were kept frozen (−40°C) in a nitrogen atmosphere prior to analysis. Samples from the outflowing centrifuged water (10 l) were filtered through pre-combusted (4 h at 450°C) glass fiber filters (Whatman GF/F, 47 mm, 0.7 µm nominal pore size) in order to check for the efficiency of the centrifugation process concerning to total solid, organic matter and lipid retention. Total suspended solids and organic matter content was gravimetrically determined after drying (60°C) filter-retained material until constant weight and subsequent combustion at 450°C, respectively. Glass fiber filtered water samples were used for nutrient analysis (nitrate, nitrite, ammonia, silicate and phosphate), all of which were spectrophotometrically carried out according to standard methods [24]. Data on rainfall and irradiance for the studied period were obtained from the nearby meteorological station of Isla Mayor, included in the Andalusia environmental network information system for agriculture managed by IFAPA (http://www.juntadeandalucia.es/agriculturaypesca/ifapa/ria/servlet/FrontController?action=Static&url=listadoEstaciones.jsp&c_prov). Data on daily Alcala Dam discharge was retrieved from the Confederación Hidrografica del Guadalquivir database, located at http://www.chguadalquivir.es/saih/Informes.aspx.

Total lipid extraction was performed on subsamples between 0.8 g and 2.6 g from the lyophilized seston material depending on the total inorganic particle content. Lipids from the dried material were extracted following the method of Folch [25] using three subsequent extractions of a 2:1 chloroform-methanol mixture. Extractions were combined into a single volume and phase separation achieved after adding a 20% volume of a 1 M KCl solution. The upper phase was discarded and the lower phase was evaporated under a stream of nitrogen. The dried total lipid extract was gravimetrically determined and stored in 2 ml Teflon sealed glass vials at a concentration of 10 mg ml^−1^ in a 2:1 chloroform/ methanol mixture containing 0.01% hydroxy-butyl-toluene (BHT). For fatty acid analysis, lipid extracts were trans methylated in an acid catalysed reaction for 16 h at 50°C using 1 mL of toluene and 2 ml of 1% sulphuric acid (v/v) in methanol [26]. Fatty acid methyl esters were separated in n-hexane and quantified in a Shimadzu GC 2010-Plus gas chromatograph equipped with a flame-ionization detector (280°C) and a fused silica capillary column Suprawax-280 (15 m × 0.1 mm). Hydrogen was used as the carrier gas. The initial oven temperature was kept at 100°C for 0.5 min, followed by an increased rate of 20°C min^−1^ until achieving a final temperature of 250°C that was kept for 8 additional min. Most individual phytoplankton fatty acid methyl esters (FAME) were identified by reference to authentic standards (Sigma 18919 C4-C24 FAME mix 18919 and Sigma 47085 PUFA Nº3 from menhaden oil). Identification of 16:4n-3 was achieved thanks to the use of a *Chlamydomonas reinhardtii* lipid extract and those diatom specific FAs (16:2n-4, 16:2n-7, 16:3n-4 and 16:4n-1) used to compute the specific PU16D diatom marker were identified using previously well-characterized FAME mixes obtained from pure cultures of the diatoms *Chaetoceros gracilis* and *Phaeodactylum tricornutum* [27]. Results were expressed as percent fatty acid respecting to the total fatty acid content since the vast majority of published FAs reference data for taxonomic ascription is in that unit [11].

### Models used to infer phytoplankton community structure

A multivariable matrix with 268 rows of seston samples (two or three replicates for each of the four seston samples taken in each of the 25 sampling times) and 20 columns representing fatty acid biomarkers (S1 Table) was used as input data in an adaptation of the CHEMTAX analysis [14] for its use with FAs instead of pigments. Previous works [12,18] had validated the usefulness of CHEMTAX when operating with FAs as biomarkers. The reference data matrix was produced from updated information on fatty acid composition recently compiled [11] for a set of 18 different marine and freshwater taxonomic phytoplankton groups. The close fatty acid profile similarity between classes in the pairs Chlorophyceae-Trebouxiophyceae, Pyramimonadophyceae-Mamiellophyceae and Coccolithophyceae-Pelagophyceae, recommended considering such pairs as single taxonomic groups to get a clearer output from the analytical model. The reference data matrix was therefore constituted by the 3 former pairs, the classes Cyanophyceae, Chlorodendrophyceae, Cryptophyceae, Pavlovophyceae, Dinophyceae, Eustigmatophyceae, Raphidophyceae, Euglenophyceae, Chrysophyceae, Xanthophyceae, Synurophyceae and the phylum Bacillariophyta, which fatty acid ratios to 16:0 are indicated in S2 Table. As in [18], 16:0 was selected as the unit fatty acid because of its highest abundance and frequency among microalgae. Since CHEMTAX uses factor analysis and a steepest descent algorithm to determine the best fit of the input data matrix to the reference matrix, 60 different runs were performed on 60 randomized ratio matrices [28]. Only those 6 outputs with the lowest Root Mean Square (RMS) error were computed for estimation of taxonomic groups contribution to the estuary phytoplankton community structure.

The seston fatty acid matrix for all GRE samples (S1 Table) was also used to infer phytoplankton community structure from the Bayesian mixing model Fatty Acid Source Tracking Algorithm in R (FASTAR), adapted by Galloway and colleagues [29] from the stable isotope mixing model MixSIR [30]. The reference library input matrix utilized consisted of the mean and standard deviation of fatty acids in the formerly described 15 microalgae groups (S2 Table). To calculate the proportional contribution of possible phytoplankton groups to the community the model iteratively assesses potential combinations of the possible phytoplankton groups to select combinations that best reflect the FA profiles found in the community sample. The “id” code for each mixture sample was treated as a fixed effect, estimating the contribution of each group to each sample separately (sample size of 1). No trophic fractionation was used (discrimination matrix set to 0) since dietary signatures were not the goal. Results of the model describe the probability of the proportional contribution by each phytoplankton group to the community in each sample. The posterior distributions were estimated using the Gibbs sampling algorithm of Markov Chain Monte Carlo (MCMC) implemented using the open-source Just Another Gibbs Sampler (JAGS) software [31]. Following previously published mixing models, the analysis implemented a normal likelihood function. MCMC chains were run as “very long” for 10^6^ iterations with 5*10^5^ iteration burn-in and a thinning rate of 5*10^2^.

### Statistical analysis

Quantitative data resulting from CHEMTAX analysis on the relative contribution of the 15 taxonomic groups to phytoplankton community structure were analysed using permutational analysis of variance included in the PERMANOVA+ add-on package for PRIMER v6 [32]. A two-way crossed factor (salinity and season) design was employed in a PERMANOVA analysis using 9999 unique permutations and significant terms were tested using a-posteriori pairwise comparisons with the PERMANOVA t statistic. Canonical analysis of principal coordinates (CAP) [33] was also performed on phytoplankton assemblages, with the analysis constrained both for salinity and season in order to find that axis through the multivariate cloud that is best at separating the a priori established groups. The first squared canonical correlation (δ_1_^2^) and the leave-one-out allocation success were used to indicate how well groups were discriminated by the CAP model. Both parameters provided an estimation of the misclassification error and showed between groups differences in the multivariate space. The relationship between environmental variables and phytoplankton community structure was investigated using a distance-based linear model (DistLM) [32] performed on their similarity matrix, with data variation partitioned according to a multiple regression model. To quantify the variance in the response variable explained by all explanatory variables, a stepwise (forward and backward) procedure and the Akaike information criterion (AIC) [34] were employed as the model selection criteria. Seasonal differences within taxonomic groups due to estuary salinity were tested using a one-way ANOVA of the arcsine-transformed data, followed by a Tukey post-hoc test.

## Results

### Efficiency of the continuous centrifugation process

Due to the infrequent application of continuous centrifugation to collect suspended particles in estuarine waters and the use of a yet untested centrifuge for such a purpose, some measurements were performed in order to check efficiency of the centrifuging process. Quantification of total suspended solids (TSS), particulate organic matter (POM) and total lipids (TL) in the outflowing water from the centrifuge revealed retentions of 95.7±1.3 (TSS), 92.3±2.7 (POM) and 92.9±1.8 (TL) within the centrifuge rotor. Increasing centrifugation speed to 12000 g was tested in some of the samples, but no significant enhancement in particle retention was found. The fatty acid profile from total lipids in the outflowing water averaged for seven randomly distributed samples differed from that in the centrifuged material (S3 Table). The nearly seven percent lipid eluding centrifugation was particularly enriched in 18:0 and 18:1n-9 whereas FAs of 20 carbons chain and longer were not detected. Other specific FAs were less represented in the outflowing collected lipids with the only exception for the slightly higher value, although nonsignificant (Anova one-way, P>0.05), detected in the sum of bacterial fatty acids (BFA, S3 Table).

### Taxonomic identification of estuarine microalgae under light microscopy

Despite visual difficulties, direct microscopic observation of estuarine water samples allowed identifying 53 microalgae genera belonging to 11 of the 18 taxonomic groups available in the reference fatty acid library. Table 1 shows the relative cell abundance after compiling by season and salinity range those genera belonging to the same class. More detailed information on phytoplankton composition at the genus level is provided in S1 Fig. Results on relative cell abundance were produced in order to get a parallel qualitative estimation of phytoplankton dynamics respecting to results obtained from FAs analysis. The Bacillariophyta was the group more frequently detected, representing over 50% of total cell abundance in most instances, with minimum contribution to phytoplankton community (26.6-28.8%) during summer in the lower salinity sector (0-10 ppt) of the estuary (Table 1). Both centric and pennate diatoms were similarly represented in the estuary (S1 Fig). Small coccoid and filamentous Cyanophyceae represented the second most abundant microalgae cells, being most frequent in summer within the 0-10 ppt salinity range (Table 1). During all seasons, there was a clear decrease in Cyanophyceae cell abundance as salinity increased. A similar, although less marked, tendency with regard to salinity was observed for the Chlorophyceae. The presence of this class was, however, more constant among seasons in comparison to the Cyanophyceae. Dinophyceae and Coccolithophyceae species were more frequently observed in winter but their pattern of variation relative to salinity was uncertain (Table 1). All other classes included in Table 1 were intermittently detected and represented less than 5% of total cell abundance.

**Table 1:**
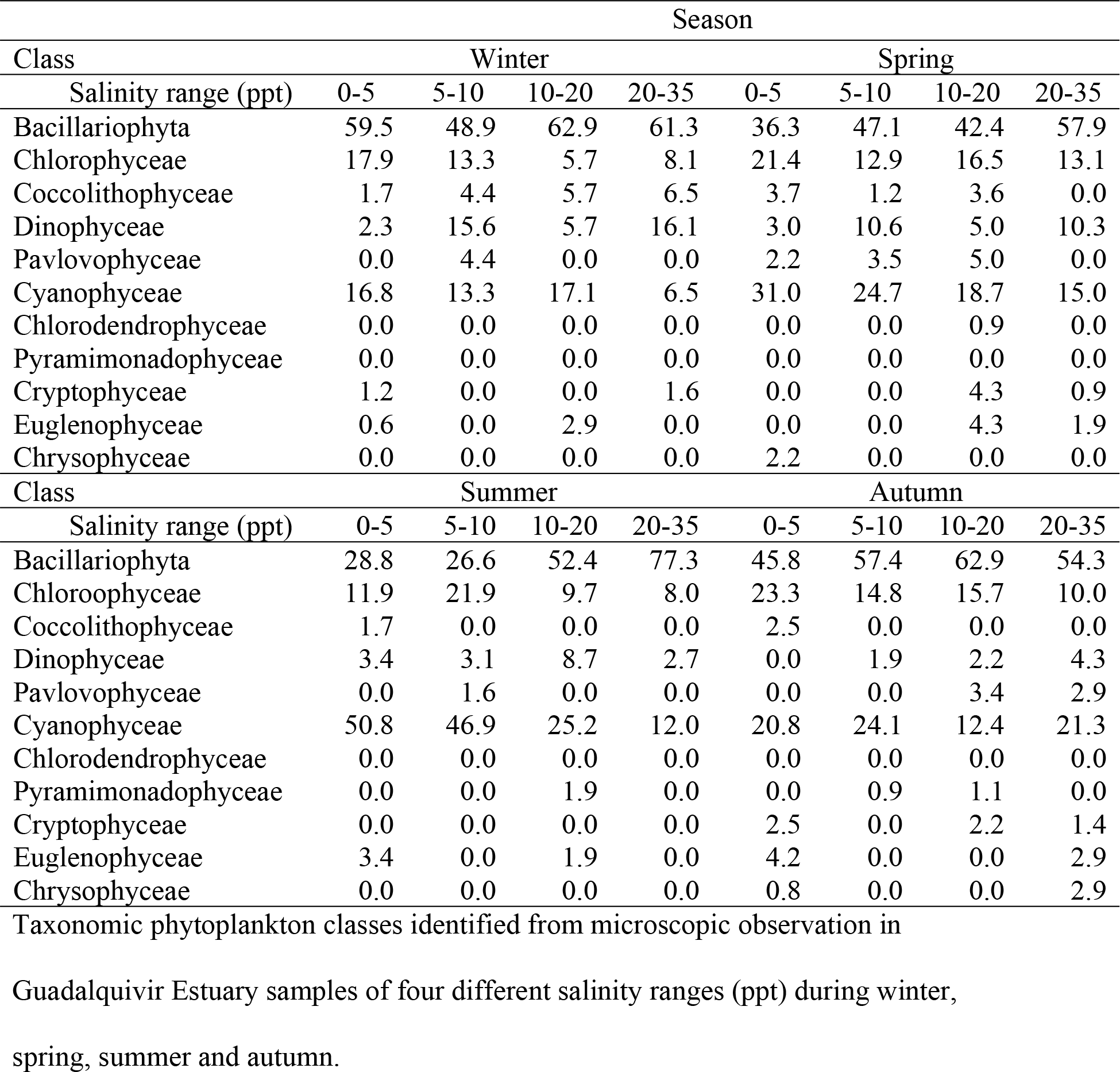
Relative abundance of phytoplankton taxonomic groups.

Phytoplankton diversity was determined for each season and salinity combination after grouping the identified taxa at the genus level. The standard Shannon-Weaver diversity index significantly increased (P<0.05) in the higher salinity sector of the estuary but only during winter and summer (Fig 1). When phytoplankton diversity was measured utilizing the higher taxonomic rank functional groups that resulted after CHEMTAX analysis, the Shannon-Weaver index decreased (P<0.05) only at the highest salinity range during spring and winter (Fig 1).

**Fig 1.**
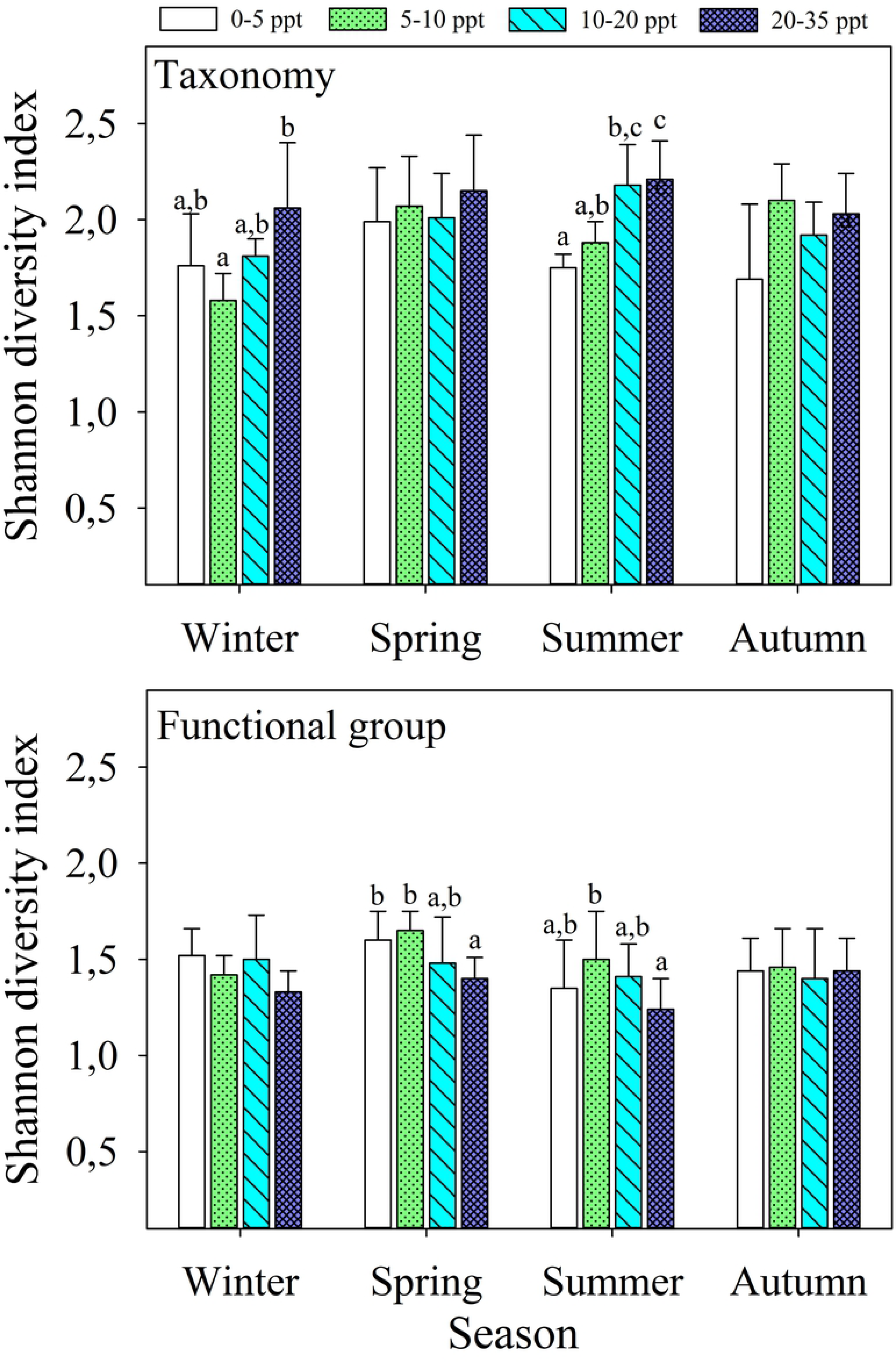
Diversity of the phytoplankton community. Shannon-Weaver diversity index were calculated from taxonomy at the genus level (top graph) and from CHEMTAX functional groups (bottom graph) for the phytoplankton community of the GRE at four different salinity ranges during Winter, Spring, Summer and Autumn. Different low case letters denote heterogeneous (P<0.05) salinity groups within each season.

### Spatiotemporal changes of phytoplankton groups inferred from fatty acids

Results of the CHEMTAX outcome indicated Bacillariophyta was the most abundant group contributing rather constantly around 50% to total phytoplankton community spatial and temporally, with only a slight, although significant (P<0.05), decrease in spring (Fig 2). The microscopically observed inverse relationship between Cyanophyceae and salinity coincided with results from CHEMTAX analysis (Fig 2). Total Cyanophyceae contribution to phytoplankton community estimated from FAs was lower than that microscopically determined. A similar conclusion can be reached for the Chlorophyceae+Trebouxiophyceae group, although with a lower difference between both methods. FAs indicated increased Dinophyceae contribution to phytoplankton community at higher salinity during spring, summer and autumn. A similar positive relation with salinity was found for Coccolithophyceae+Pelagophyceae abundance (Fig 2). The presence of Eustigmatophyceae representatives was caught by CHEMTAX analysis whereas species from this class could not be detected under the microscope. The Eustigmatophyceae contributed around 10% to phytoplankton community in winter, spring and summer, and around 5% in autumn, but no evident relationship with salinity could be concluded (Fig 2). Despite significant seasonal effect in the contribution of the six more represented taxonomic groups in the phytoplankton community, the overall results illustrated in Fig 2 (with maybe the exception of the Dinophyceae) point to the permanent presence of these phytoplankton groups throughout the year in the GRE.

**Fig 2.**
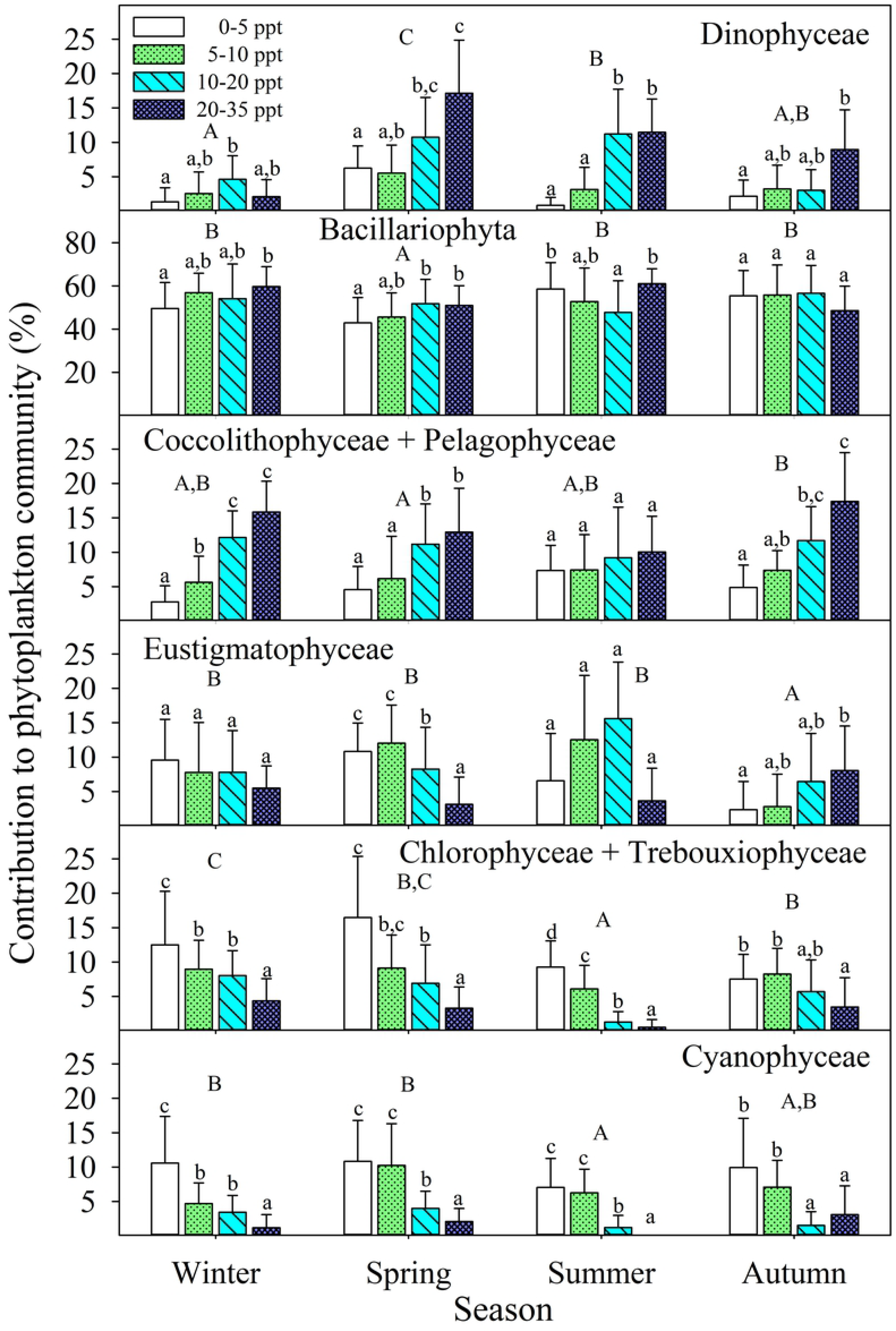
Most represented phytoplankton groups from CHEMTAX analysis. Percent contribution to community of phytoplankton classes based on their fatty acids. Homogeneous salinity groups within each season are denoted by the same low case letter and differences among seasons are depicted with capital letters.

Among the minority taxonomic groups, occurrence of the Raphidophyceae was only perceived after CHEMTAX analysis. It was shown to be the most marked seasonal group (detected only in spring and summer) and it was present only in the higher salinity waters of the estuary (Fig 3). The Euglenophyceae showed an inverse trend, being noticeably more abundant in the autumn-winter period and in lower salinity waters. The Pavlovophyceae was also more abundant during autumn-winter but it favourably proliferated in higher salinity waters (Fig 3). Other interesting seasonal change was the near absence of Pyramimonadophyceae, Mamiellophyceae and Cryptophyceae representatives during winter. The Chlorodendrophyceae was the most seasonally constant group (Fig 3). The classes Chrysophyceae, Xanthophyceae and Synurophyceae were not detected by any of the matrix factor analysis, coinciding with their practical absence in microscopic observations (Table 1).

**Fig 3.**
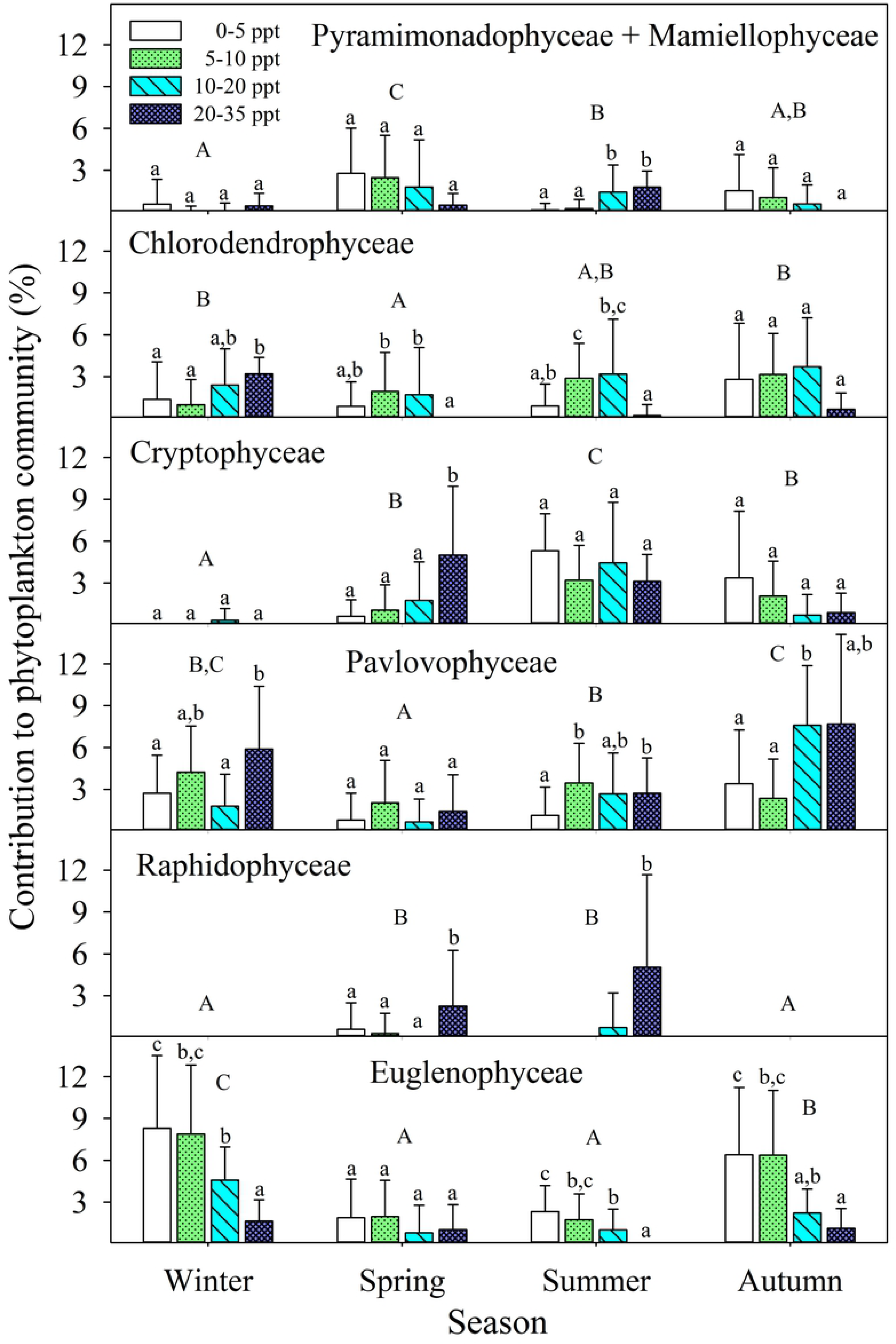
Least represented phytoplankton groups from CHEMTAX analysis. Percent contribution to community of phytoplankton classes based on their fatty acids. Homogeneous salinity groups within each season are denoted by the same low case letter and differences among seasons are depicted with capital letters.

Comparing CHEMTAX results with those inferred by the FASTAR Bayesian analysis indicated an overall similar group contribution to phytoplankton community, but some subtle differences existed depending on the inferring method. Dispersion of results was more frequently elevated following the Bayesian analysis, as indicated by higher standard deviations in most of the abundant taxa (S2 Fig) and, particularly, in the least represented taxa (S3 Fig). The Bacillariophyta was also the most represented group following FASTAR, however its percent contribution to community was generally lower respecting to CHEMTAX results. The spatiotemporal distribution pattern of the Dinophyceae was similar regardless the inferring method. There was, nevertheless, a single between-method relevant mismatching point for the highest salinity in winter for which CHEMTAX inferred very low Dinophyceae contribution (Fig 2) whereas FASTAR estimated the highest contribution (S2 Fig). The higher percent contribution calculated by CHEMTAX for the Coccolithophyceae-Pelagophyceae and Eustigmatophyceae groups contrasted with the lower values produced by FASTAR, a circumstance that was inverse to that for the Cyanophyceae (S2 Fig). The Eustigmatophyceae was practically absent in the phytoplankton community during autumn-winter according to FASTAR and similarly distributed temporally according to CHEMTAX results. Such disagreement could not be contrasted with microscopy due to the impossibility of identifying small coccoid microalgae commonly included in the Eustigmatophyceae. Among the less represented phytoplankton groups the most remarkable fact was the nearly double percent contribution to community estimated for the Euglenophyceae following FASTAR analysis (S3 Fig) in comparison to CHEMTAX results. The seasonality of this class was, on the other hand, similarly estimated by both inferring methods. Chlorodendrophyceae, Cryptophyceae and Pavlovophyceae were less represented in the phytoplankton community when FASTAR was employed (S3 Fig).

### Environmental drivers of phytoplankton community structure dynamics

The permutational analysis of variance in the multivariate space demonstrated highly significant effects for season and salinity as well as for the interaction between both factors (Table 2). Salinity explained more percent of total variation than season but the highest source of variation (36.62%) was residual (Table 2). The CAP performed on the full data set of samples revealed a noticeably more structured pattern across axis 1 (δ_1_^2^=0.705, δ_2_^2^=0.032) when constraining was done on salinity (Fig 4). In this instance, samples from lower to higher salinity clearly scored from left to right on axis 1 with minimal variation through axis 2. The Coccolithophyceae+Pelagophyceae was the main phytoplankton group positively correlating with axis 1, whereas Cyanophyceae and Chlorophyceae+Trebouxiophyceae were the most important groups showing negative correlation with axis 1. Constraining the CAP model on season resulted in a more disperse sample ordination, with a weaker first squared canonical correlation (δ_1_^2^=0.405) and a higher δ_2_^2^ (0.299). Samples from the autumn-winter period principally scored on the left side of axis 1 and the spring-summer samples principally scored on the right side (Fig 4). The important source of variation recovered by axis 2 reflected the salinity gradient within seasons. The stronger influence of salinity on sample ordination was evidenced after CAP constraining on season since, even in this case, higher salinity samples scored separately from lower salinity samples. Under such ordination pattern, Coccolithophyceae+Pelagophyceae, Dinophyceae, Raphidophyceae and Cryptophyceae were the phytoplankton groups more related to higher estuary salinity waters, and Chlorophyceae+Trebouxiophyceae, Cyanophyceae and Euglenophyceae the groups more related to lower salinity (Fig 4). Separating seasonal and salinity influence on sample ordination revealed the salinity gradient exerted a weaker effect on structuring phytoplankton community in spring (δ_1_^2^=0.548) while higher δ_2_^2^ value (0.375) in summer reflected elevated inter annual variation in this season (Fig 5) that coincided with maximal inter annual contribution to variability calculated by PERMANOVA in spring (Table 3). Winter was the season showing the lowest inter annual variation.

**Table 2.**
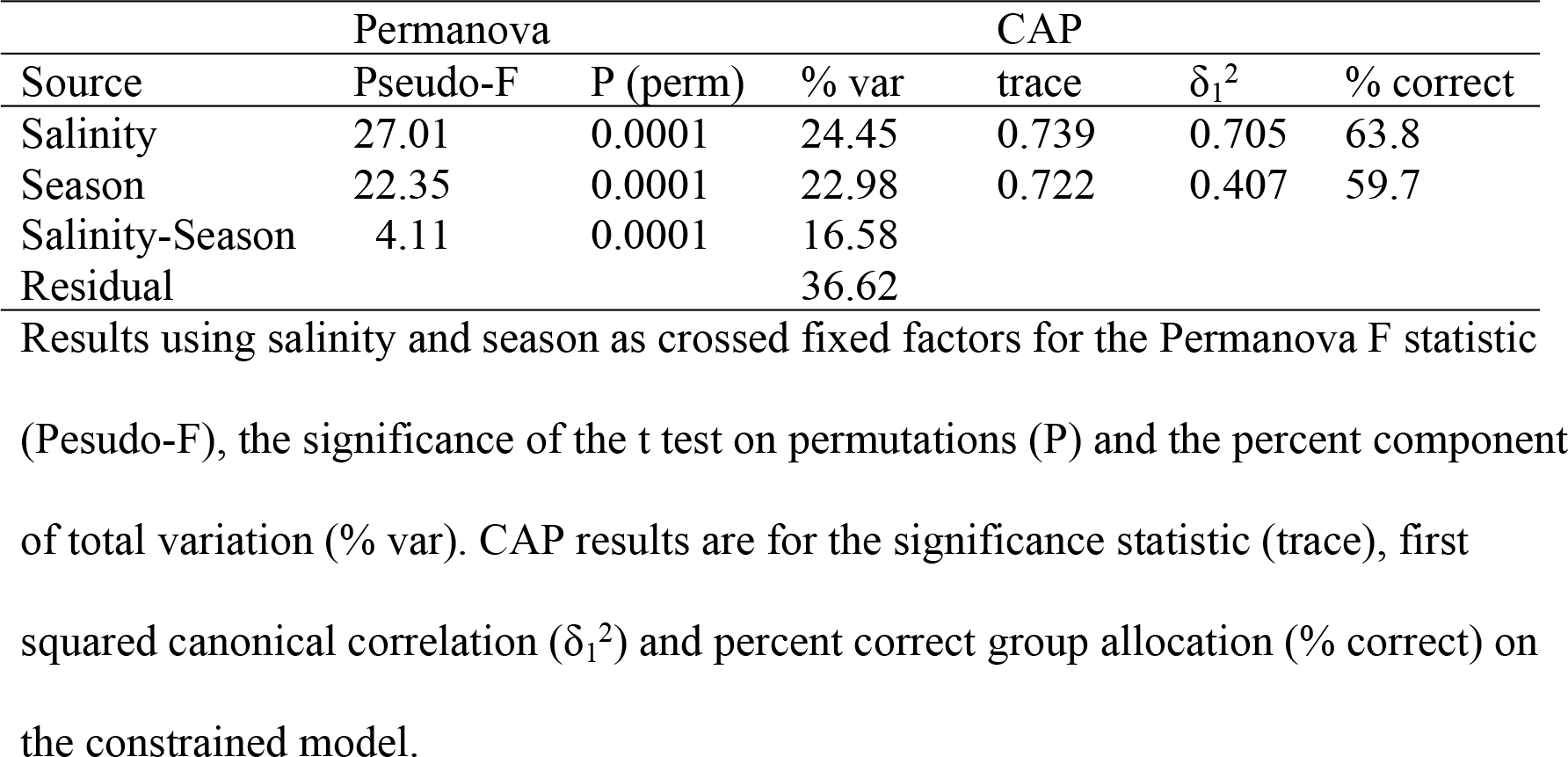
Output of the Permanova and canonical analysis of principal coordinates (CAP) on the phytoplankton community structure.

|  | Permanova |  |  | CAP |  |  |
| --- | --- | --- | --- | --- | --- | --- |
| Source | Pseudo-F | P (perm) | % var | trace | $\delta_1^2$ | % correct |
| Salinity | 27.01 | 0.0001 | 24.45 | 0.739 | 0.705 | 63.8 |
| Season | 22.35 | 0.0001 | 22.98 | 0.722 | 0.407 | 59.7 |
| Salinity-Season | 4.11 | 0.0001 | 16.58 |  |  |  |
| Residual |  |  | 36.62 |  |  |  |

Results using salinity and season as crossed fixed factors for the Permanova F statistic (Pseudo-F), the significance of the t test on permutations (P) and the percent component of total variation (% var). CAP results are for the significance statistic (trace), first squared canonical correlation ( $\delta_1^2$ ) and percent correct group allocation (% correct) on the constrained model.

**Fig 4.**
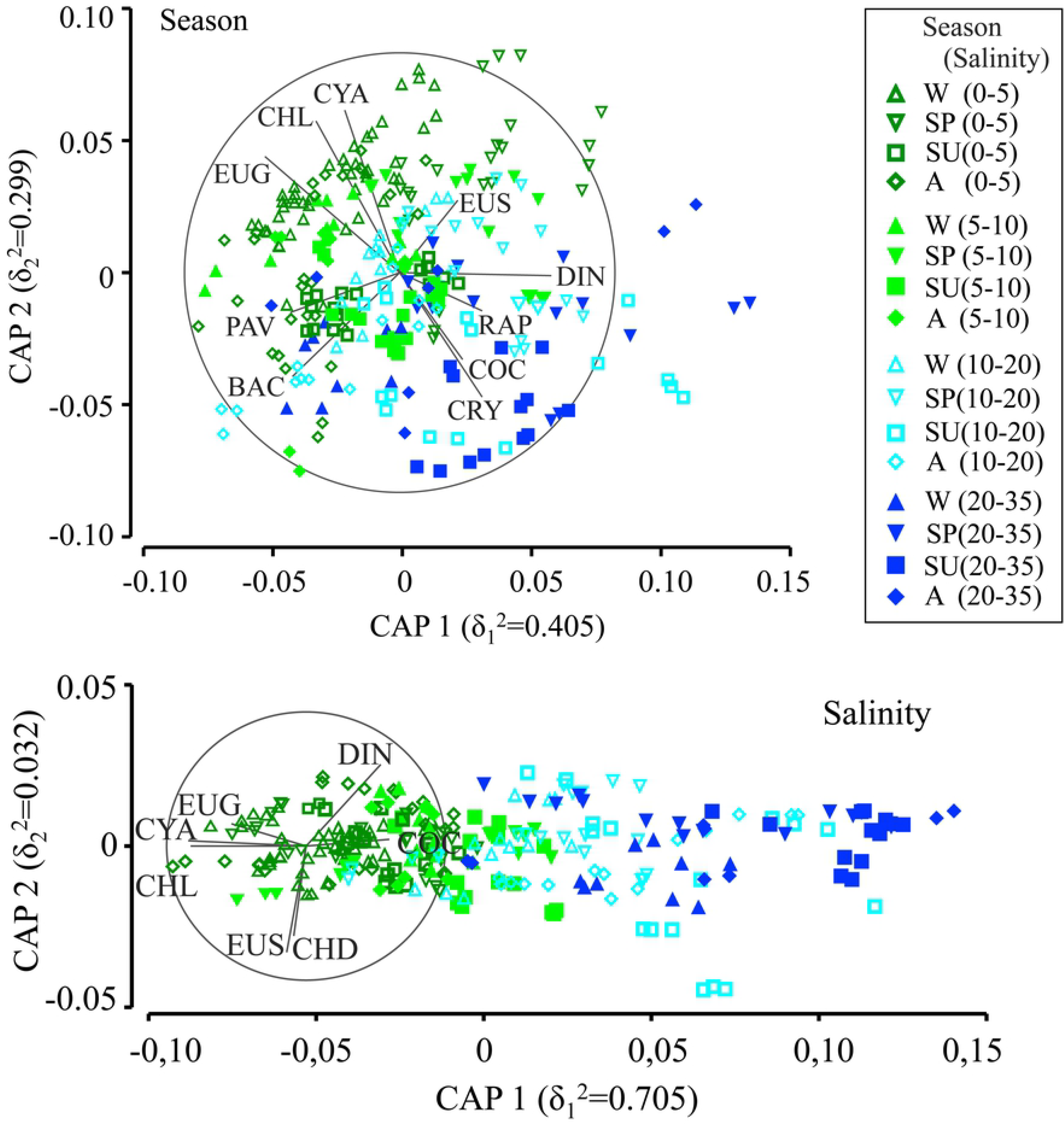
Annual sample ordination after canonical analysis of principal coordinates (CAP) on the phytoplankton community structure. The relative contribution of phytoplankton taxonomic groups inferred from fatty acids was used as descriptor variables in models constrained either by season (top graph) or salinity (lower graph).

**Fig 5.**
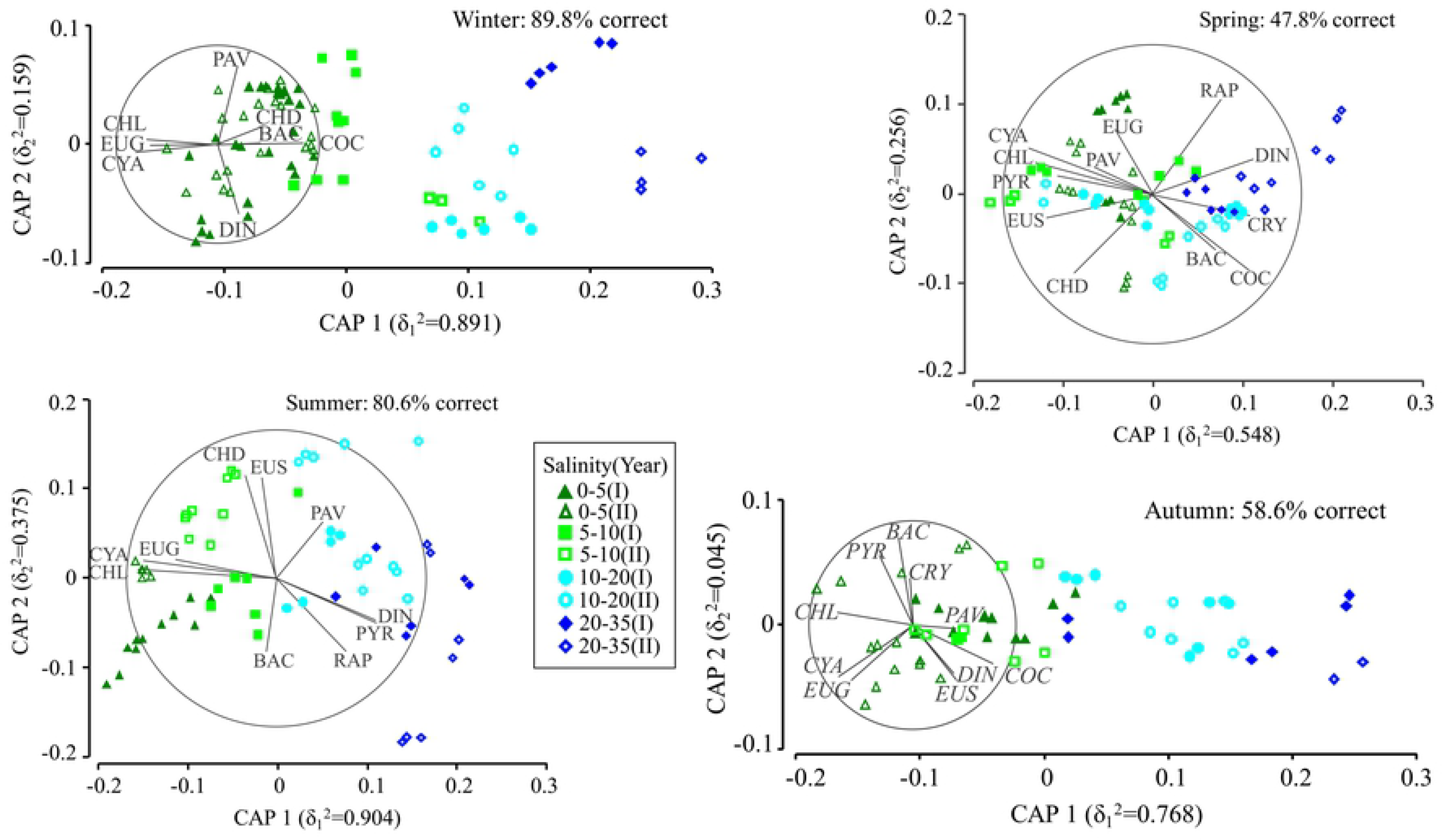
Seasonal sample ordination after canonical analysis of principal coordinates (CAP) on the phytoplankton community structure. The relative contribution of phytoplankton taxonomic groups inferred from fatty acids was used as descriptor variables in a model constrained by salinity (four levels of variation). Inter annual variation within each season can be noticed from differences between year I (solid symbols) and year II (empty symbols).

**Table 3.** Salinity and inter annual percent components of total variation in the phytoplankton community structure.

|  | Season |  |  |  |
| --- | --- | --- | --- | --- |
| Source | Winter | Spring | Summer | Autumn |
| Salinity | 32.64 | 27.12 | 29.87 | 27.58 |
| Year | 15.13 | 22.39 | 29.87 | 25.99 |
| Salinity-Year | 21.74 | 20.26 | 13.54 | 9.58 |
| Residual | 30.47 | 30.21 | 26.71 | 36.88 |

Percent source contribution obtained after performing two way crossed fixed factors
Permanova during winter, spring, summer and autumn.

In a preliminary DistLM analysis, nutrients were included among all environmental predictor variables and were found to explain low percent variation of phytoplankton composition, namely 1.4% (phosphate), 1.8% (ammonia), 4.4% (silicate) and 5.6% (nitrate). Low nutrient contribution to total variation and their very high levels detected throughout the estuary salinity gradient (Fig 6), discouraged using nutrients as relevant explanatory variables and were thus removed from the analysis. Despite nitrate, phosphate and silicate concentrations being significantly lower in higher salinity waters, their respective minimum values were still too high to be considered as potentially limiting factors of phytoplankton growth [35]. Ammonia showed an inverse trend respecting to other nutrients, increasing with salinity in the estuary, and its relative contribution to the inorganic nitrogen pool was nearly an order of magnitude lower than that of nitrate (Fig 6). Nitrite concentrations were on average half the value of ammonia and no clear spatial or temporal pattern could be found. Nutrient concentrations were unaffected by dam discharge (Fig 7). When seasonality was decomposed into the five environmental predictor variables included in Fig 8, DistLM revealed salinity explained most of the phytoplankton community structure variation in winter, spring and summer. Rainfall was the most important factor in autumn, coinciding with the typically elevated rainfall variability occurring in the zone, but having minor influence during the dryer summer and winter periods (Fig 8). The contribution of dam discharge to explain phytoplankton community variability was very constant at around 10-11% from spring to autumn and reached a minimum 2% in winter. The effect of temperature was more marked in summer-autumn respecting to the minimal influence exerted in winter-spring. The relevance of irradiance was maximal in summer and reached the minimum value in autumn (Fig 8).

**Fig 6.**
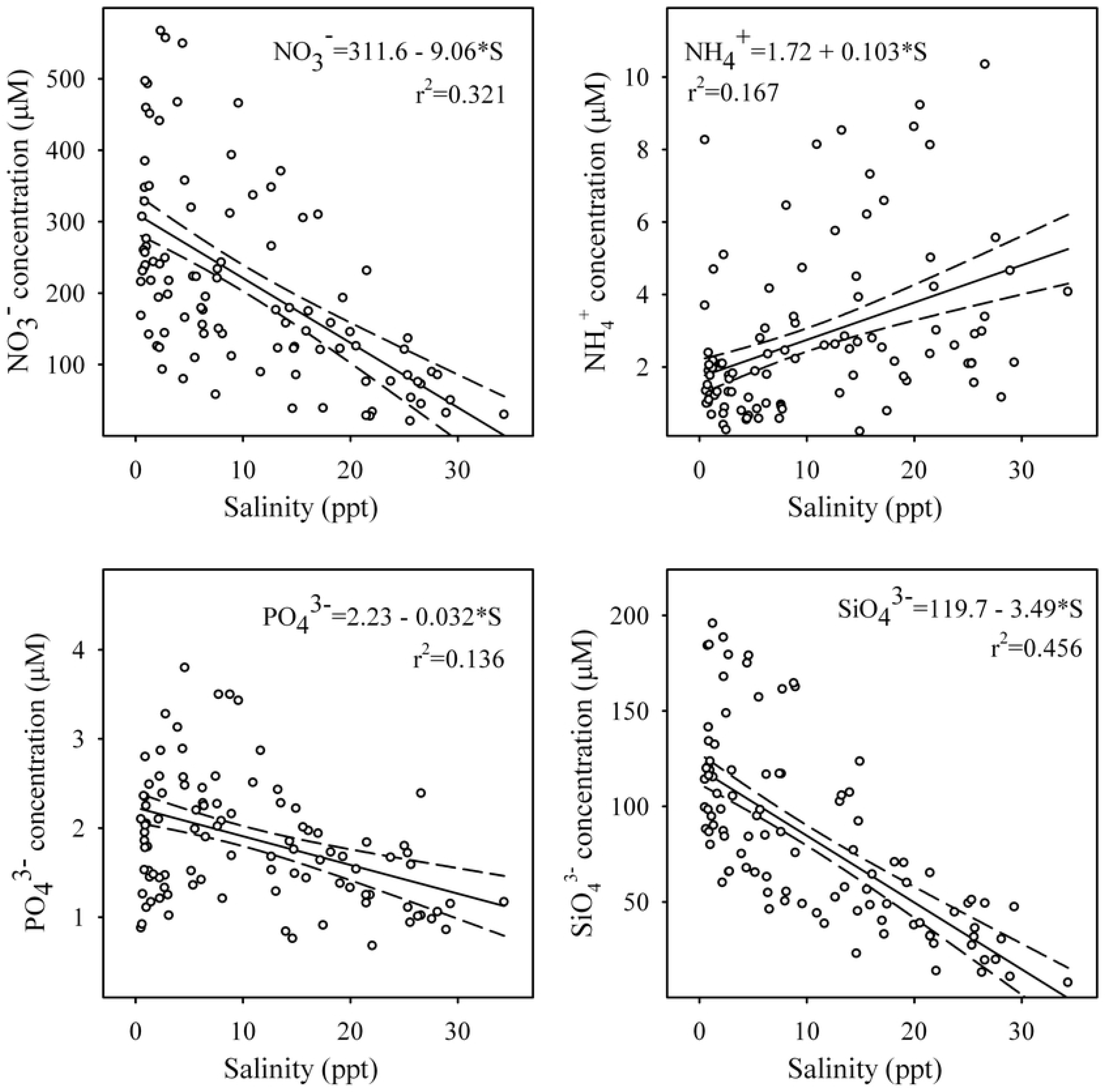
Relationship between nutrients and salinity in the Guadalquivir estuary. Concentrations (µM) of nitrate, ammonia, phosphate and silicate are regressed against salinity throughout the annual period.

**Fig 7.**
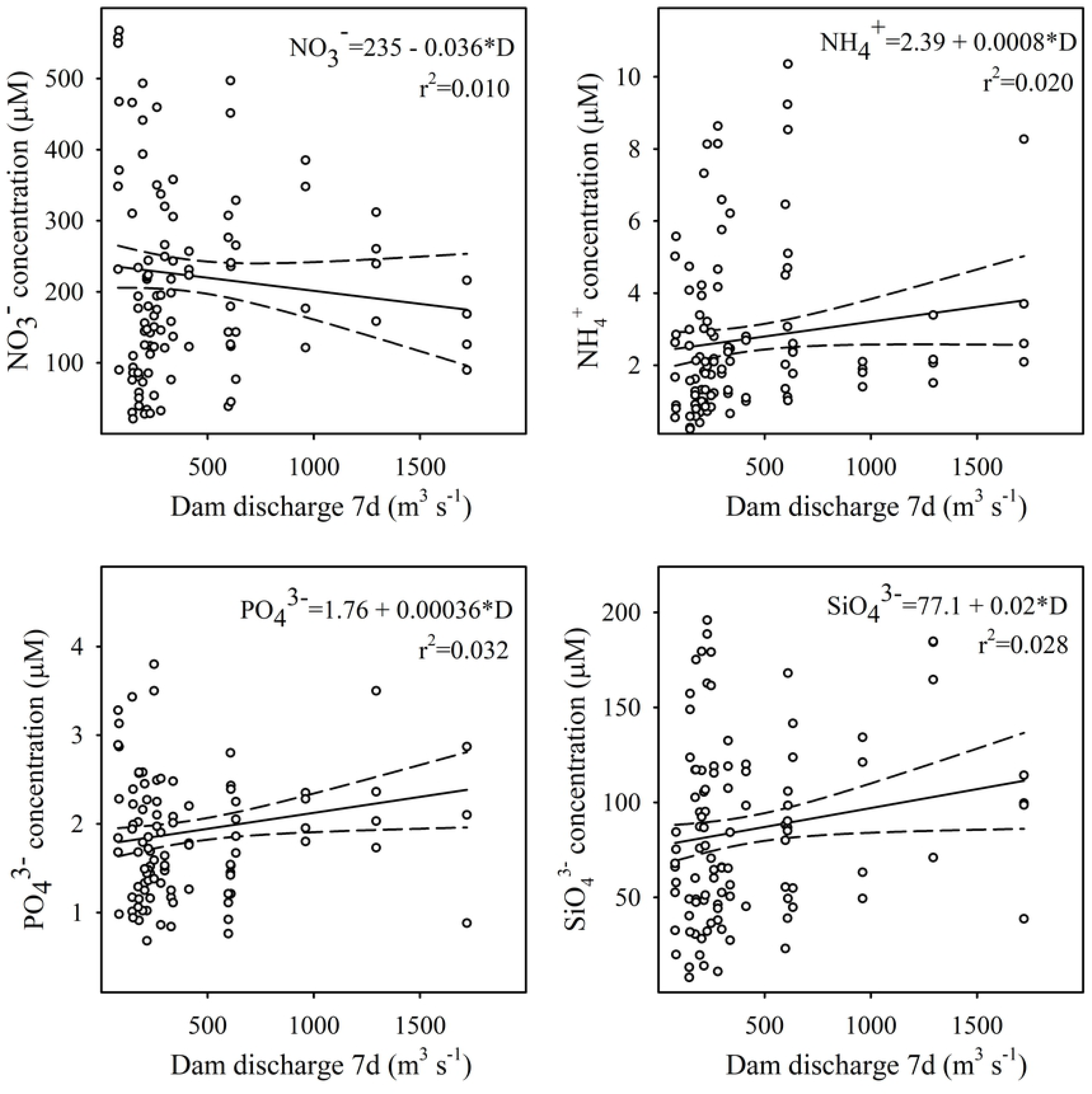
Relationship between nutrients and the Alcala dam discharge in the Guadalquivir estuary. Concentrations (µM) of nitrate, ammonia, phosphate and silicate are regressed against dam discharge throughout the annual period.

**Fig 8.**
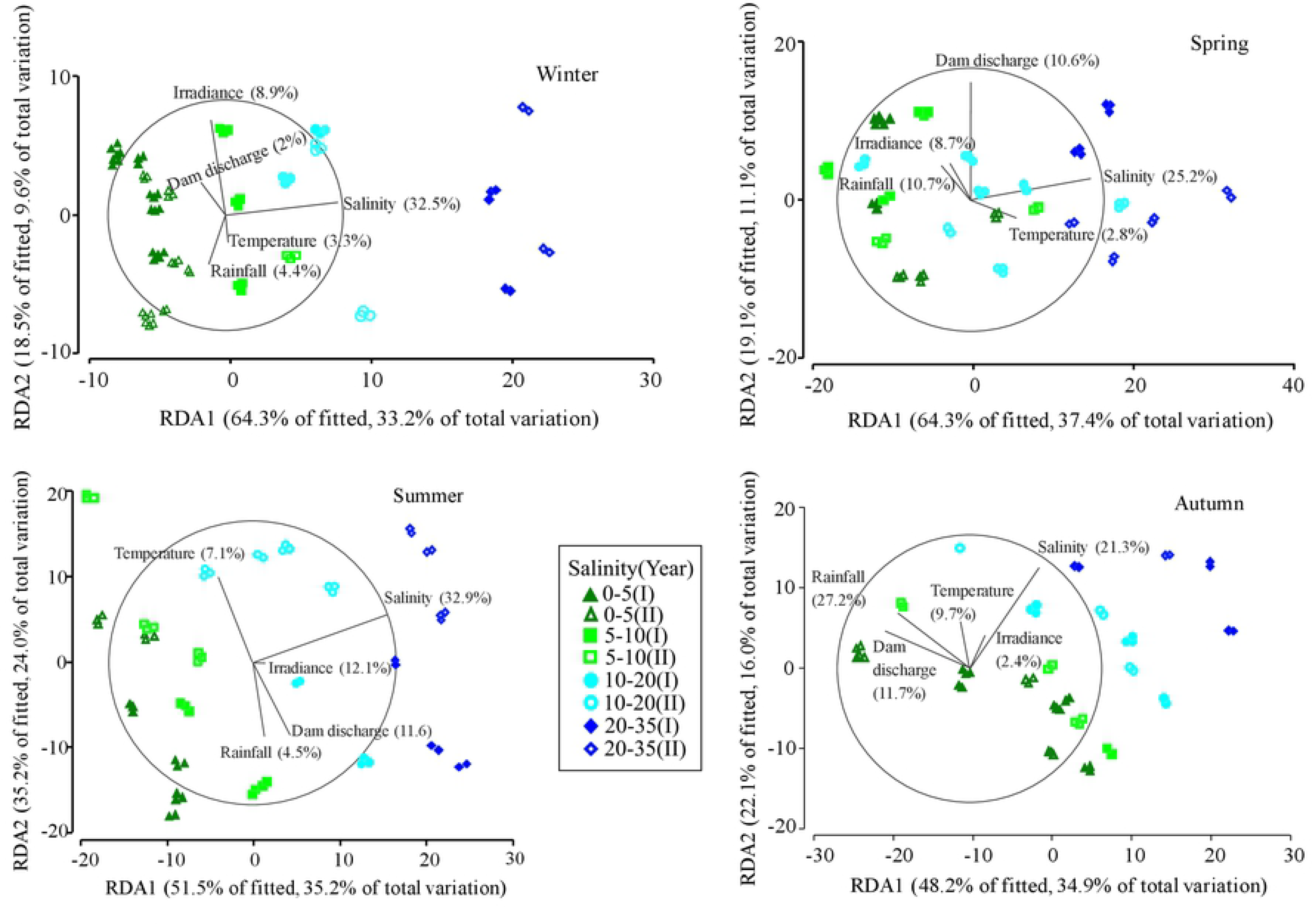
Distance-based redundancy analysis (dbRDA) of the phytoplankton community using the complete set of predictor variables. The percent contribution of the predictor variables salinity, temperature, irradiance, rainfall and dam discharge to variability of total phytoplankton community (denoted in parenthesis) was calculated for each season. Inter annual variation within seasons can be noticed from differences between year I (solid symbols) and year II (empty symbols).

## Discussion

### Suitability of the fatty acid based procedure to quantify structure of phytoplankton assemblages

The combined use of continuous centrifugation of estuarine water for seston collection, its subsequent FAs determination and the novel implementation of matrix factor analysis produced the largest quantitative estimation of any aquatic phytoplankton community structure known to date. Continuous centrifugation of GRE water was demonstrated to efficiently collect TSS in water samples that are difficult to process using conventional filtration. Although the application of continuous centrifugation in estuarine phytoplankton research has been very little referred in the literature, it cannot be considered a novel concentration procedure since an early work [36] already described its usefulness and particularly high efficiency in estuarine waters. The high centrifuge retention efficiency for TSS in the GRE is similar to values recently achieved for riverine samples employing a simple, on-site usable and easy-to-operate continuous flow centrifuge [37].

The elevated proportion of inert material in centrifuged and freeze-dried seston samples did not represent any analytical restriction for lipid extraction since they were processed as normally done with sediment samples [38]. Entering seston FAs as input data in both CHEMTAX and FASTAR resulted in unprecedented quantitative seasonal and spatial information on the relative abundance for 12 out of the 18 taxonomic groups available in the reference library. Within the 12 groups identified using FAs, those of the Eustigmatophyceae and Raphidophyceae can be remarked as the ones that could not be detected from microscopy. There are only three previous published works on the use of FAs to quantify the structure of phytoplankton communities and they resolved five phytoplankton groups in the Schelde Estuary [12], eight different classes in Finnish Lakes [19] and three phytoplankton groups in combination with fungi and bacteria in temporary ponds [18]. Contrary to present study, the classes Chlorophyceae and Trebouxiophyceae could be differentiated in the Schelde Estuary [12]. On the other hand, Dinoflagellates and Haptophytes could not be discerned, as has been achieved in present work. FAs-based inferences of present work enabled including taxonomic contribution to community of small nano and picophytoplankton species which have been traditionally enumerated just as unidentified coccoid or flagellate components of the community by conventional microscopy. FAs-based analysis is therefore capable of revealing the presence of an increased number of phytoplankton functional groups that have been routinely neglected by microscopy [2]. Since phylogeny optimally represents functional differences among taxa [39], it seems evident that ecosystems functioning can be better understood when a more exhaustive description of their constituents is achieved. Pigment based analysis has also contributed to enhance taxonomic determination of phytoplankton communities [40–42] and their combined application with FAs has generated very interesting synergic results [12, 43].

Although quantifying differences between FAs-based and microscopy methods was not the goal of present study, the general coincidence in the estimation of phytoplankton group contribution to community determined from both approaches in those taxonomic groups that could be identified under the microscope can be remarked. Similar results have been found in other works using FAs as chemo markers [19, 44, 45]. Current information at this respect is nevertheless still very limited to allow achieving any robust conclusion. From reviewed similar works using pigment as biomarkers [46], it can be learnt that there is agreement [47–50] but also disagreement [7, 51, 52] for results contrasting pigment CHEMTAX analysis and microscopy. More studies are thus required to confirm whether this pigment-related observed variability could also occur when FAs are used as biomarkers. With current knowledge, it can be thought that FAs based inferring could show less variability than pigment based studies thanks to the more elevated influence of taxonomy on phytoplankton FAs respecting to environmental effects [10]. Compared to pigments, intraspecific changes in FAs due to irradiance are minimal [12]. It has also been shown that fatty acid profile variability due to phosphorus stress is lower than that associated to phylogeny [27].

Despite subtle differences between CHEMTAX and FASTAR outcomes, their similar overall quantification of phytoplankton community structure is in agreement with both inferring methods producing similar estimations [53]. Lower standard deviations following CHEMTAX analysis can be related to the process of randomization and selection of the 10% outputs with the lower RMS error recommended for optimal operation of CHEMTAX [28]. Contrary to CHEMTAX, the FASTAR analysis utilizes standard deviation values in the reference matrix. In this regard, the currently available information on phytoplankton FAs reflects elevate data dispersion within functional groups [11] that could have induced increased standard deviations in the estimated composition matrix [53]. The lower percent contribution to community inferred from CHEMTAX for Dinophyceae in winter and Cyanophyceae in autumn were two remarkable disagreements respecting to FASTAR results that deserves further studies. The higher contribution inferred by CHEMTAX analysis for Coccolithophyceae+Pelagophyceae, Pavlovophyceae and Eustigmatophyceae could not be compared with microscopy results due to the unfeasibility for identifying species of these taxa in the GRE waters. However, some important representatives (*Isochrysis* and *Diacronema*) of these groups were found to bloom [54] and were isolated and cultured from ponds receiving only GRE water [55].

The lower relative contribution to community of Cyanophyceae and Chlorophyceae+Trebouxiophyceae determined from FAs inference in comparison to microscopy results was likely associated to the inability of transforming cell abundance into biomass of the phytoplankton functional groups. The evident difficulty to achieve reliable cell abundance to biomass conversion factors for several coccoid, small cell-sized and string or colony forming species of these two groups, is of particular severity in the highly turbid waters of the GRE. Such inherent complexity for adequate phytoplankton visualization in turbid estuaries make it necessary to develop new tools based on taxa-specific chemo markers. Advantages at this regard can be deduced from present results on the Eustigmatophyceae. Although species of this class are commonly coccoid, their fatty acid profile is well differentiated and that making it possible that both CHEMTAX and FASTAR could have detected this class which is not feasible by microscopy. Pigment analysis revealed Eustigmatophyceae species are common in estuarine waters [56]. Pico-coccoid microalgae have been found to represent up to 93% of total phytoplankton cell volume under certain conditions in some estuaries while cyanobacteria can reach 6% of total biovolume [42]. The Raphidophyceae was other class exclusively detected when FAs were used as biomarkers, and its contribution to phytoplankton community was low and restricted to higher salinities. It could be that a very low cell density could have preclude microscopy identification of Raphidophyceae species. They are found in estuaries but could not be detected when pigments were used as biomarkers likely due to absence of specific reference pigments [7]. This represents other example on the synergic effects that can be achieved when pigments and FAs are used in combination as phytoplankton biomarkers. CHEMTAX and microscopy results for Cryptophyceae suggests contribution of this class to phytoplankton community in the GRE is negligible, contrasting with higher expected results for a class that shows particular eco-physiological adaptations to estuarine waters [57].

### Spatiotemporal dynamics of phytoplankton groups inferred from seston FAs

Despite the intrinsic difficulty for microscopic visualization of microalgae species in the highly turbid GRE waters, a set of 53 different genera could be identified without including those unidentified coccoid and microflagellate microalgae. A similar result has been reported in managed ponds receiving exclusively GRE waters [54]. If the generally accepted underestimation of phytoplankton species richness using microscopy [58] is also assumed, it is obvious the actual number of phytoplankton taxa must be noticeably higher in the GRE. Phytoplankton richness in the GRE can be comparable to that of other estuaries describing 97 taxa (belonging to a total of 44 genera) in the Schelde Estuary [59], 45 genera in South Africa St. Lucia Estuary [60], 55 genera in the Kowie Estuary [61] and those indicating the presence of 63 genera following a molecular approach in the Vire River Estuary [62]. A similar molecular approach detected up to 101 genera in the Segura river estuary [63]. The estimation of phytoplankton diversity at the genus taxonomic rank in this study can explain the slightly lower mean indices here obtained respecting to other works using the species level [64]. When diversity was determined on the functional groups resulting after CHEMTAX analysis, the mean Shannon-Weaver index around 1.5 was very close to the central value of the diversity range (0.77-2.29) calculated from photo-pigments in Galveston Bay [65]. It is also worth considering that present diversity index range of variation (1.08-1.84) was smaller, with higher minimum values, than those reported in similar works [64, 65]. Such stability and moderately high phytoplankton diversity in the GRE would correspond to an ecosystem characterized by elevated resource use efficiency and stable functions [66]. Ecosystem stability could be the consequence of specific conditions concerning uninterrupted nutrients sufficiency, river flow hyper-regulation preventing typical seasonal washout events [20] and an elevated permanent turbidity discouraging peaks of primary production. Taxonomic diversity provided useful complementary information to results on community structure achieved after FA-based inferring methods. Computing phytoplankton diversity on the basis of a higher taxonomic rank than that of species (genus from microscopy or class from chemo-markers) is important in functional diversity to achieve a better grasp on ecological processes [67].

Traditional studies based on qualitative FAs indices [68] contributed advancing knowledge on the qualitative structure of phytoplankton assemblages in estuaries [15–17, 43] and other aquatic ecosystems [13, 44, 69–71]. Results here presented on the quantitative determination of twelve phytoplankton groups during the full year cycle and for the complete salinity gradient of the GRE have no precedent in any other estuary. It has been evidenced the use of FAs as taxonomic biomarkers represents a new step forward to advance in studies similar to those focusing on seasonal changes in estuarine phytoplankton community composition along the complete salinity gradient [52, 59, 72–74]. The elevate spatial stability of phytoplankton diversity in the GRE has also been described throughout a river-estuary continuum [61]. In present study, such stability implied that no diversity maximum occurred in the 5-10 ppt salinity range, as has been estimated from reviewed information in the Baltic Sea [75]. However, a recent high throughput molecular study could not detect any maximum or minimum of phytoplankton diversity in the low salinity transition reaches of the Baltic Sea [74], in consonance with present results for the GRE. With the exception of diatoms, the succession among main functional groups along the salinity gradient revealed by CAP analysis of the CHEMTAX outcome is consistent with the concept of ecocline described at the species level in the Schelde Estuary [59]. Temporal predominance of diatoms has also been observed in other estuaries subjected to different environmental conditions [60, 72, 76–78]. It has also been pointed out that in turbid and eutrophic estuaries, diatoms dominate the phytoplankton community throughout the year [79]. Lack of any detected diatom bloom in the GRE was likely related to high and permanent turbidity and contrasts with the more marked diatom seasonality and typical blooms occurring in other temperate region estuaries experimenting higher environmental fluctuations [80]. The presence of diatoms throughout the complete salinity gradient in the GRE is also consistent with reports for other nine Western Europe estuaries [72].

The decline in Chlorophyceae+Trebouxiophyceae quantitative contribution to phytoplankton community along the GRE salinity gradient coincides with estimations from pigment analysis studies reporting Chlorophytes dramatic decreasing from freshwater to salinities of 15-24 ppt [76]. Chlorophytes also decreased considerably from the lowest salinity to a value around 20 ppt in the Segura River Estuary [63]. Similarly, Chlorophytes dominated (40-60%) phytoplankton community in the freshwater reaches of the Wankan River Estuary and represented less than 4% when salinity reached 10-15 ppt [52]. Although the Chlorophyceae+Trebouxiophyceae did not show a marked seasonality in the GRE, FAs indicated their significant lower presence during summer, contrasting with the late spring-summer peak reported for the Nervion Estuary [73]. Low seasonal variability was also inferred from FAs in the Cyanophyceae, with lack of any typical summer peak that has also been found in marsh ponds receiving waters from the GRE [54, 55]. This is an opposite situation to that described in the Neus River Estuary [3] and the less turbid and less nitrogen loaded Guadiana Estuary where Cyanobacteria consistently exhibited summer peaks [21]. Contrary to seasonality, salinity exerted a strong influence on Cyanophyceae abundance and the gradual decline observed with increased salinity is in accordance with observations for the Tagus Estuary [76], the Wankan Estuary [52] and the phosphorus limited Mississippi and Pearl River Estuary [81]. The increasing abundance of Coccolithophyceae+Pelagophyceae with salinity revealed by FAs analysis is consistent with the markedly marine nature of the Coccolithophyceae [82] and the frequent presence of this group in the higher salinity estuarine reaches [63, 73]. The ability to differentiate between the dynamics of two haptophytes (Coccolithophyceae and Pavlovophyceae) using FAs represents a novel finding respecting to other biochemical procedures. The slightly more elevated presence of the Pavlovophyceae during autumn-summer in the GRE cannot be compared to other estuaries due to lack of reference information. However, the presence of the Pavlovophyceae in the GRE can be deduced from the finding that *Diacronema* is a common blooming species in the fringing marsh ponds [54, 55].

Proliferation of the Dinophyceae in higher salinity waters of the GRE and during the warmest period (spring-summer) agrees with results in other estuaries [76, 83]. The low abundance of the Cryptophyceae in the well-mixed GRE contrasts with the high relevance of this class in the stratified waters of the fjordic Loch Linnhe [50] and also with their increased importance associated to salinity and during summer-autumn in the Tagus Estuary [76]. The relatively stable presence of the Eustigmatophyceae inferred from FAs analysis along the GRE salinity gradient has not been reported in any other estuary. Euglenophyceae proliferation during the autumn-winter period in the GRE could represent a slightly temporal displacement respecting to the Tagus Estuary where this class is more abundant during summer-autumn and it is almost absent the rest of the year [76]. A decreased presence of the Euglenophyceae along the salinity gradient has also been observed in the Segura River Estuary [63]. Among the less represented classes in the GRE, the small changes in the Chlorodendrophyceae, Pyramimonadophyceae and Mamiellophyceae associated to seasonality and salinity represent the first report on their quantitative contribution to phytoplankton community. Because Prasinophytes are generally difficult to identify [84], little is known about their abundance, although the application of molecular tools enabled identification of the genus *Pyramimonas* as the exclusive Chlorophyte representative in the highest salinity reaches of the Segura River Estuary [63]. As inferred from FAs in the GRE, the Raphidophyceae was also found to have a minor contribution to phytoplankton community in the Nervion Estuary [73]. Its preferential presence in the higher salinity waters and during the warmest season is consistent with results in other estuaries [83, 85].

### Factors driving changes in phytoplankton community structure

The multivariate analysis of FAs-inferred spatiotemporal changes in phytoplankton community structure revealed salinity as the main influencing factor. The CAP analysis suggested that salinity effects surpassed those other factors related to seasonality. Salinity has been acknowledged to play a primary role in structuring estuarine phytoplankton communities and canonical multivariate analysis has determined its main role in different estuaries [61, 86]. Nevertheless, salinity was not found to exert such influence in the Pearl River Estuary [87], exemplifying the elevated variability in the response of estuarine phytoplankton community to spatial changes generally remarked [1]. The strong influence of salinity on phytoplankton community structure in the GRE is likely related with the attenuation of other influencing environmental factors due to anthropogenic activities. That would be the case of decreased fluctuations in hydrology (minimization of high wash-out and drought events) due to river damming, chronic low light availability due to persistent turbidity and apparently unlimited nutrient resources. Temperature (range 10.2°C-30.0°C) was obviously less affected than the above factors by anthropogenic activities and it explained little of phytoplankton community variability in the GRE, contrasting with strong influence of this factor in the Neuse River Estuary, where temperature range (6.7°C-29.3°C) was characterized by lower minimum values [3]. It is, nevertheless, complicated to ascertain whether this difference between estuaries can be mostly attributed to temperature regime since its interaction with other environmental factors can notably differ between both estuaries. In this regard, the average suspended solids and the nitrogen to phosphorus ratio in the GRE are around 25-fold and 10-fold higher than the respective mean values reported for the Neuse River Estuary [3]. The permanently elevated suspended solids in the GRE probably influenced solar irradiance to explain little of phytoplankton community variability.

The persisting high nutrients content in the GRE minimized their influence on phytoplankton community structure respecting to other environmental factors. Although references using a similar canonical multivariate assessment to that here used are very limited, some works similarly point to low nutrient effects in estuaries. In this regard, nutrients had also limited effect on phytoplankton community in the Tagus Estuary [86] and played a secondary role in the Baltic Sea [88], the Pearl River Estuary [87], the Neretva River Estuary [88] and even in the oligotrophic Adriatic coast exposed to direct influence of the Po River [90]. On the other hand, ammonia emerged as an important regulating factor in the Neuse River Estuary [3], highlighting the importance of specific cases within the diversified estuarine phytoplankton response to nutrients. In general terms, nutrients can be regarded to influence estuarine phytoplankton in a lower extent than that found in open coastal systems [45]. River flow is other important component of seasonality that in the GRE is under the influence of the marked raining periods in spring and autumn, typical of the Mediterranean climate, and the strong dam human control that attenuates the effects of the naturally occurring rainfall. The exceptional 27.2% contribution of rainfall to total variation of phytoplankton community in autumn, surpassing the influence of salinity during this season, was the most remarkable fact concerning the effects of river flow and coincided with the heaviest and more variable rainfall recorded during the studied period. It also illustrates the limits of the Alcala Dam to regulate river flow during peaking rainfall, something expected from the relatively low water impoundment capacity of the dam respecting to volume of the estuary [20]. With the exception of autumn, the low relevance of rainfall on phytoplankton community structure in other seasons is consistent with elevated residence time (40-140 d, for 80% of the studied period) of the river discharge in the GRE, estimated from dam discharge data and estuary volumetric values already reported [23]. In the GRE, river flow seems to be a weaker force at structuring phytoplankton community compared to other estuaries [76, 91].

## Conclusions

The setting up of a procedure for collecting particulate matter through continuous flow centrifugation in a highly turbid estuary allowed determining FAs that were successfully applied to quantitatively infer phytoplankton community structure. Entering seston FAs data in matrix factor analysis (CHEMTAX or FASTAR) utilizing the largest known to date reference FAs library provided unprecedented information on the spatiotemporal dynamics of twelve taxonomic groups in the yet unexplored phytoplankton of the GRE. Both taxonomic and functional group diversity derived from FAs-based analysis resulted similar to diversity indices in other estuaries. All analytical methods revealed diatoms dominated spatiotemporally the phytoplankton community in the GRE. Matrix factor analysis also indicated the abundance of Cyanobacteria, Chlorophytes and Euglenophytes was inversely related to salinity, whereas an opposite relationship occurred for the Coccolithophyceae, Pelagophyceae and Dinophyceae classes. FAs revealed seasonality was more marked in some of the less represented groups (Raphidophyceae, Euglenophyceae) while seasonal changes were moderate in the most abundant groups of the phytoplankton community. Human induced stability and/or saturation in some environmental factors (river flow, turbidity, eutrophication) positively contributed to seasonal stability in the phytoplankton community.

## Supporting information

**S1 Fig. Relative cell abundance after microscopic identification of phytoplankton taxonomic groups in the GRE.** Species abundance of all identified taxonomic groups in water samples obtained within salinity ranges of 0-5, 5-10, 10-20 and 20-35ppt during winter, spring, summer and autumn.

**S2 Fig. Most represented phytoplankton groups from FASTAR analysis.** Percent contribution to community of phytoplankton classes based on their fatty acids. Homogeneous salinity groups within each season are denoted by the same low case letter and differences among seasons are depicted with capital letters.

**S3 Fig. Least represented phytoplankton groups from FASTAR analysis.** Percent contribution to community of phytoplankton classes based on their fatty acids. Homogeneous salinity groups within each season are denoted by the same low case letter and differences among seasons are depicted with capital letters.

**S1 Table. Seston fatty acids in the Guadalquivir Estuary.** Mean fatty acid values of seston samples from four different salinity ranges during Winter, Spring, Summer and Autumn.

**S2 Table. Reference fatty acid ratio matrices.** Mean fatty acid ratios to 16:0 reference matrix used in the CHEMTAX analysis (S2.1) and mean±standard deviation reference matrix used in FASTAR analysis (S2.2) to infer phytoplankton community structure in GRE seston samples. The output ratios from CHEMTAX analysis are shown in S2.3.

**S3 Table. Efficiency of the centrifuging process.** Mean and standard deviation of fatty acid profile (percent of total fatty acids) from GRE samples centrifuged in a continuous centrifuge and their corresponding particulate matter collected in the outflowing water flow.

## Acknowledgments

Authors are deeply grateful to Carmen Pérez Gavilán and Alejandra Ramirez de Arellano for their assistance in sample processing and fatty acid analysis.

## Author contributions

**Conceptualization:** José-Pedro Cañavate, César Vílas.

**Data curation:** José-Pedro Cañavate, Stefanie van Bergeijk.

**Formal analysis:** Stefanie van Bergeijk, Enrique González-Ortegón.

**Funding acquisition:** César Vílas.

**Investigation:** José-Pedro Cañavate, Stefanie van Bergeijk, César Vílas.

**Methodology:** José-Pedro Cañavate, Stefanie van Bergeijk, Enrique González-Ortegón.

**Project administration:** César Vílas.

**Supervision:** José-Pedro Cañavate, César Vílas.

**Writing – review & editing:** José-Pedro Cañavate.

